# Replication of treatment effects and differences among populations during experimental evolution of sex allocation in an annual plant

**DOI:** 10.1101/2024.12.12.628098

**Authors:** Nora Villamil, Xinji Li, John R. Pannell

## Abstract

Variation among replicates of experimental evolution studies that begin by sampling from a single source population can be attributed to measurement error or to random differences in the environment experienced by measured individuals. On the basis of measurements in a large common garden, Cossard et al. (2021: *Current Biology* 7, 1-7) reported rapid evolution of enhanced ‘leaky’ sex allocation by females (which produced male flowers) in experimental populations of the dioecious herb *Mercurialis annua* from which males had been removed. Their study thus demonstrated the rapid dissolution of dioecy via changes in sex allocation in response to mate limitation and strong competition for siring success. But their study also found substantial variation among replicate populations. Here, we replicated their common garden, growing plants from the same populations but under different conditions and assaying the measured variables differently. The effects (significance, magnitude and direction) of selection treatment and generation on the sex expression of females with inconstant sex expression met five definitions for successful replication. Importantly, comparisons between the two studies revealed relatively consistent differences among the replicate populations, suggesting that the among-population variation reported by Cossard et al. cannot be attributed to random noise or local differences in growing conditions in the common garden. Our study represents a rare example of the replication of a common garden study and raises interesting questions about the nature of interpopulation divergence within treatment groups of studies using experimental evolution.

## INTRODUCTION

The rate of adaptive evolution in a population depends on the strength of natural selection and the narrow-sense heritability of the trait on which selection is acting (Falconer and Mackay, 1996). Because narrow-sense heritability is the additive genetic proportion of phenotypic variance for the trait in question, the rate of adaptive change depends critically on the degree to which the trait is phenotypically plastic: the more phenotypic variance is attributable to plasticity, the lower is its heritability and the rate of its evolutionary change (Falconer and Mackay, 1996). The ability of experimental evolution studies to detect evolutionary divergence over short periods of time between populations evolving under contrasting conditions thus depends, to an important extent, on reducing environmental variation within and among replicate populations for a given treatment. Similarly, assays of evolutionary change after a period of divergent evolution are designed to culture organisms in environmentally homogeneous environments, e.g., in common gardens, in which all individuals from all populations experience much the same conditions.

Experimental evolution typically involves subjecting populations to one of two or more different environments (treatments) over a number of generations, with several replicate populations assigned to each treatment. Of most interest is usually the treatment effect, which is detected in terms of divergence in the average phenotype of individuals in populations under one treatment from that under the other treatment(s). As in any experiment, differences among replicate populations within a particular treatment diminishes our power to detect divergence among treatments, but such population differences may be biologically interesting in their own right too. For example, populations subjected to the same selection pressure may evolve differently because they began with different genetic variants as a result of sampling variance at the start of the experiments, because of genetic drift (Kutnjak et al., 2017), because different populations acquired different mutations over the course of the experiment (Travisano et al., 1995), or perhaps because of pre-existing plasticity in the trait under selection that leads to differential survival followed by genetic changes (Corl et al., 2018).

Cossard et al. (2021a) reported rapid divergence in sex allocation between experimental populations of the wind-pollinated dioecious annual plant *Mercurialis annua*. As in many dioecious plants, males and females of *M. annua* are ‘leaky’ or inconstant in their sex expression, occasionally producing a few flowers of the opposite sex. Cossard et al. (2021a) wished to establish whether selection on sex allocation could bring about an evolutionary transition from dioecy to hermaphroditism. At the beginning of their experiment, they established six populations of 210 individuals each. Three were allowed to evolve as controls with a 1:1 ratio of males to females, as found in natural populations, whereas the other populations comprised only females. Over the course of four generations, females growing in the absence of males rapidly evolved a substantial increase in their degree of ‘leakiness’, underpinning a transition from separate to combined sexes. Cossard et al. (2021a) empirically showed that such reversion can occur rapidly when one sex (males) are lost, e.g., through a population bottleneck. Indeed, the evolutionary change observed in the experiment corresponded to one of the most extreme rates of divergence yet recorded in any experimental evolution study. While Cossard et al. (2021a) focused their analysis and interpretation on the overall increase in male-flower production by females in the three populations of their experiment lacking males, they also recorded substantial variation in male-flower production among the replicate populations. This variation was substantial, despite the fact that all individuals were grown together in a single common garden after the four generations of evolution.

Here, we replicate the common garden of Cossard et al. (2021a) by growing individuals from all six treatment and control populations for two of the same generations sampled in the original study. Our primary aim was to determine whether the variation in sex allocation observed among replicate populations under the same experimental treatment by Cossard et al (2021) reflects genetic divergence among these populations or whether, rather, it reflects random experimental noise. Specifically, do populations with high trait values in the previous common garden show high trait in a second common garden, or is there no co-variance in trait values between the two experiments? Although we also grew plants from the different treatments, populations and generations within a similar randomised common garden to that used by Cossard et al. (2021a), our common garden was established in a different year (with different weather), plants were grown for longer, and we used a modified sampling protocol to estimate the sex-allocation phenotypes. We used prediction intervals to determine whether differences between the studies were due only to sampling error, or due to other factors (e.g., environmental and methodological differences) (Spence and Stanley, 2016). A direct replication and comparison of the results from the two gardens allowed us to assess the robustness of those findings and establish broader grounds for generalisation of the results and their implications.

A second aim of our study was to present an example of the replication of a previous common garden experiment. Study replication provides greater confidence for generalisation, even in cases where the first results are statistically well supported and appear to be robust (Nakagawa and Parker, 2015; Parker et al., 2016; Gurevitch et al., 2018).(Kelly, 2006; Nakagawa and Parker, 2015; Parker et al., 2016; Gurevitch et al., 2018; Fraser et al., 2020). Replicate studies are rare in most disciplines (Romero, 2019; Allard and Vazire, 2021), particularly in ecology and evolution in which currently only 0.023% of studies have been replicated (Kelly, 2019). To our knowledge, ours is the first self-identified replication study of a common garden assay of treatment group and population divergence under experimental evolution.

## MATERIALS AND METHODS

### Original *versus* replicate study settings

For ten years, a long-term experimental evolution study has grown *Mercurialis annua* plants under two treatments, which we here label ‘control’ and ‘treatment’. Control populations comprised three replicate populations each with 210 individuals under a 1:1 male:female sex ratio; treatment populations comprised three replicate populations with 210 females in each. In 2016, Cossard et al. (2021a) grew seeds from all replicate populations and both treatments and from generations 1 to 4 under the same conditions in a common garden. In 2020, members from the same research group (Pannell Group, University of Lausanne) conducted a second common garden experiment in which they grew seeds from generations 1 and 3 of the experimental evolution study. Hence, this second study in 2020 replicated measures from generations 1 and 3 of the 2016 common garden experiment. Further details on plant culturing and sampling protocols are reported in the Supplementary Materials.

Despite being a direct replicate (same seed bank, season, location, facilities, same variables measured, same research group), our replication of Cossard et al. (2021) had two methodological differences. First, although plants were sown and grown following the original procedures (see Appendix 1), plants in 2020 were re-potted out of the germination trays 4 to 6 weeks later than in 2016 and were harvested at a later age, when the plants were substantially larger (greater biomass). This delay was caused by restrictions in accessing plants during the first wave of the Covid-19 pandemic. Second, the replicate sampling method consisted in subsampling male and female reproductive structures only from the top (apical) 30 cm of each plant, thus contrasting with the approach taken in the original study, where reproductive structures were sampled throughout the whole plant. Apical subsampling has potentially different consequences on male and female sex allocation estimates, because *M. annua* inflorescences are protogynous, and because fruits remain on plants longer than do male flowers male flowers, which are potentially more greatly represented towards the apex. (Villamil et al., 2021). Given such methodological differences, we expected effect sizes to differ the two common gardens, but the same patterns ought to be found with similar strengths (in terms of Cohen *d*’s scale) (Cohen, 1988). Consequently, we included a sensitivity-based approach to assess replicability (Schauer and Hedges, 2021).

### Replicability analyses

We asked whether the two main results of Cossard (2016) would be found in a new common garden: the increase in both the number of females producing male flowers; and the extent to which they did so. We also analysed the replicability of aboveground plant biomass as a housekeeping variable to compare the overall plants size differences between experiments. The replication analyses focussed on the significance and effect size (direction and magnitude) that generation, treatment and the interaction (generation x treatment) had on each response variable. Successful replication was evaluated using the five following replication definitions for each fixed effect in the model, with each response variable analysed separately:

1. Correspondence in the significance of the P-value;
2. Correspondence in the effects size direction;
3. Correspondence in the effects size magnitude (effect sizes may differ in their exact value but should fall within the same Cohen’s d scale (small: 0.2 ≥ ES < 0.5; medium: 0.5 ≤ ES < 0.8; large: ES ≥ 0.8);
4. Inclusion in the prediction interval of the original study (sensu Spence and Stanley, 2016). Due to methodological differences, we predicted our replicate underestimates male sexual function by around 10% and female sexual function by around 30% (Villamil et al., 2021). Consequently, if replicated estimates are outside the prediction interval, these are deemed to be successful replication as long as they fall 10% (♂) and 30% (♀) beyond the limits of the prediction interval.
5. Correspondence in the amount of variance explained by population identity for each response variable in both studies.

We used generalised mixed models to analyse both datasets, fitting each response variable separately within the same model structure. Our models included generation, treatment and their interaction as fixed effects. As random effects we included the population identity to account for spatial variance among sites, and population nested within generation to account for the temporal non-independence of plants within a given population but from subsequent years. All analyses were conducted using R statistical software (version 3.61) and the following R packages: lme4, MuMIn, DHARMa, predictionInterval, ggPlot, patchwork (Bates et al., 2016; Spence and Stanley, 2016; Wickham et al., 2016; Pedersen, 2017; Barton and Barton, 2018; Hartig, 2019).

### Analysis of population variation

We explored the variation among and within populations for male flower production, leakiness in sex expression, plant biomass, male reproductive effort (MRE), and male sex allocation (MSA). MRE in the original study was calculated as male flower biomass / total plant biomass; in contrast, MRE was calculated in the replicate study as male flower biomass within the subsample / subsample biomass. MSA was calculated as male flower biomass / (male flower biomass + seed biomass), again for the apical subsample. We estimated the correlation values among the population means in the original versus replicate study. Within-population variation was expressed with bidirectional error bars (SE). This analyses allowed us to assess qualitatively whether variation patterns were replicated. For each response variable modelled, we estimated the proportion of variance explained by population identity, producing a quantitative estimate. This estimate was extracted from the mixed models described above.

## RESULTS AND DISCUSSION

### Replicability of an experimental evolution study

Our repetition of the common garden experiment established by Cossard et al. (2021a) has largely confirmed the results they reported; it thus represents a successful scientific replication according to the five criteria evaluated (Figure 1, Table 1). Specifically, the two studies have yielded consistent results concerning the effects of selection treatment and generation on the proportion of females with leaky sex expression and the male flower production by females. Both the selection treatment and generation had a significant positive effect on leakiness and male flower production by females, with the selection treatment having a two-fold stronger effect on both traits (Figure 1). In contrast to these two traits, plant biomass was not a successfully replicated metric, as expected: recall that plants in the replicate study were older when harvested, and thus substantially larger.

**Table 1.** Quantitative evaluation of the five replication criteria: *P*-value significance; effect size magnitude and direction; effects size scale class; prediction interval inclusion; and percentage of variance explained by population identity. Estimates for the original (ori) and replicate study (rep) are followed by a qualitative evaluation of whether each criteria were successfully replicated (Rep?). Effect size scales are defined according to Cohen’s d scale (small: 0.2-0.49; medium: 0.5-0.79; large: ES≥0.8). The prediction interval summarises whether the mean effect size of the replicate falls within the prediction interval of the original study (i.e., whether differences are due only to sampling error). Successfully replicated parameters are shaded and ‘^*^’ indicates the result is marginally replicated.

| Response | Predictor | Study | Significance |  | Effect size |  |  | Prediction interval | Variance explained by... |  |  |  |
| --- | --- | --- | --- | --- | --- | --- | --- | --- | --- | --- | --- | --- |
|  |  |  | P-value | Rep? | direction/<br>magnitude | scale | Rep? | Within? | Group | Study | % | Rep? |
| Leakiness | Generation | ori | 0.002 | Yes | +0.30 | Small | Yes | Yes | Gen_Pop | ori | 0 | Yes |
|  |  | rep | 0.023 |  | +0.31 | Small |  |  |  | rep | 0 |  |
|  | Treatment | ori | 7.72<sup>-06</sup> | Yes | +1.02 | Large | Yes* | Yes | Pop | ori | 0 | Yes |
|  |  | rep | 0.0002 |  | +0.79 | Med |  |  |  | rep | 0 |  |
|  | Interaction | ori | 0.544 | No |  |  |  |  | Residual | ori | 66.84 | Yes |
|  |  | rep | 0.001 |  |  |  |  |  |  | rep | 71.03 |  |
|  | Generation | ori | 0.527 | Yes | +0.49 | Small | Yes | Yes | Gen_Pop | ori | 6.94<sup>-04</sup> | Yes |

| Male<br>flower<br>production |  | rep | 0.215 |  | +0.40 | Small |  |  |  | rep | 1.33<sup>09</sup> |  |
| --- | --- | --- | --- | --- | --- | --- | --- | --- | --- | --- | --- | --- |
|  | Treatment | ori | 0.003 | Yes | +0.70 | Med | Yes | Yes | Pop | ori | 4.31<sup>-11</sup> | Yes |
|  |  | rep | 0.013 |  | +0.70 | Med |  |  |  | rep | 1.18<sup>-10</sup> |  |
|  | Interaction | ori | 0.009 | Yes |  |  |  |  | Residual | ori | 0.65 | Yes |
|  |  | rep | 0.042 |  |  |  |  |  |  | rep | 2.19 |  |
| Plant<br>biomass | Generation | ori | 0.910 | Yes | +0.04 | None | No | Yes* | Gen_Pop | ori | 17.05 | No |
|  |  | rep | 0.103 |  | -0.26 | Small |  |  |  | rep | 0.58 |  |
|  | Treatment | ori | 0.292 | No | -0.28 | Small | No | No | Pop | ori | 3.53 | Yes |
|  |  | rep | 0.015 |  | -0.50 | Med |  |  |  | rep | 2.09 |  |
|  | Interaction | ori | 0.334 | No |  |  |  |  | Residual | ori | 76.44 | Yes |
|  |  | rep | 0.023 |  |  |  |  |  |  | rep | 86.44 |  |

**Figure 1.**
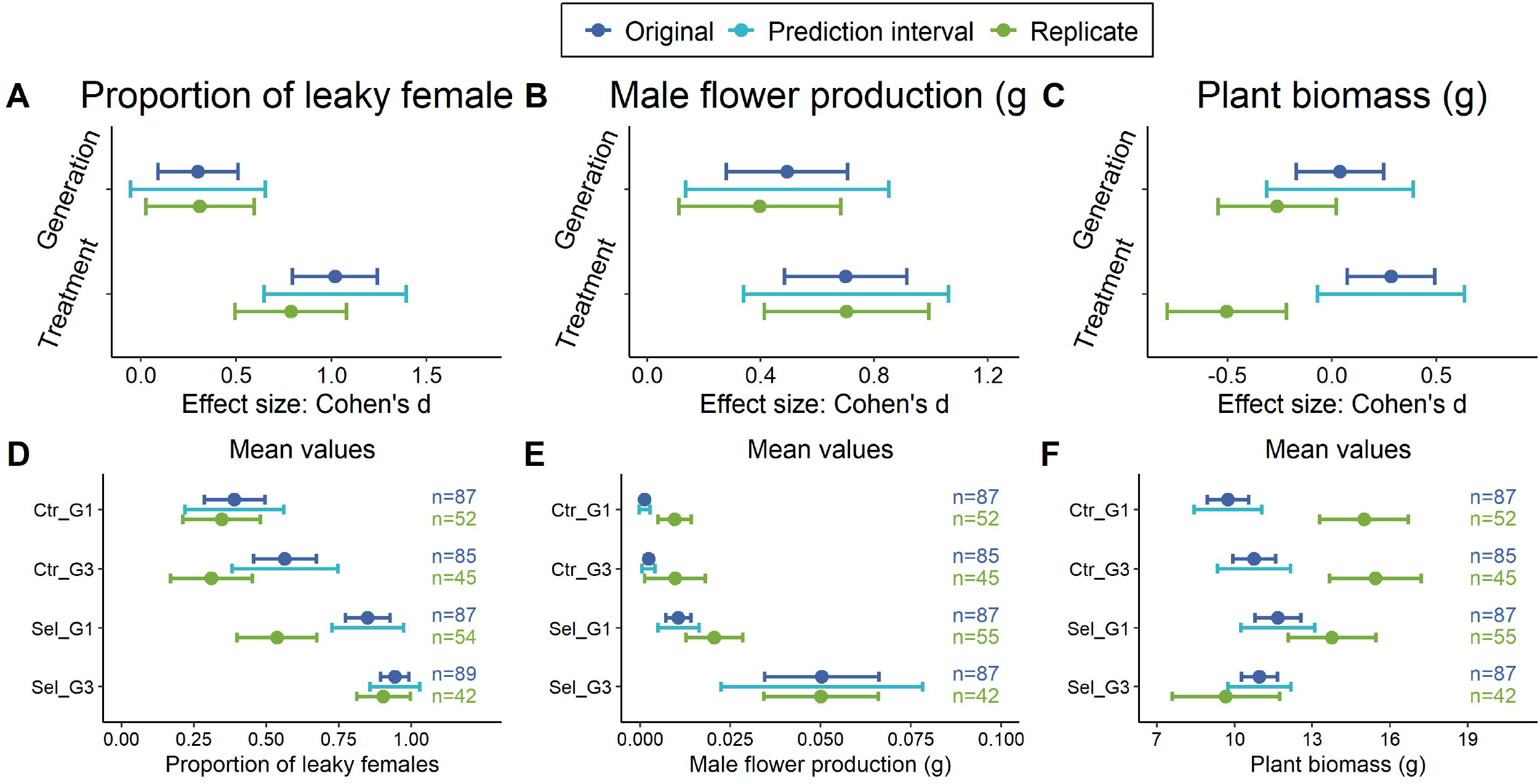
Replicability of the effect sizes (mean ± 95% CI) of generation and selection treatment of the original study (2016) *versus* those obtained in the replicate study (2020) for (A) the proportion of leaky females, (B) male flower biomass and (C) plant aboveground biomass. Replicability of the mean values per group (mean ± 95% CI) for (D) the proportion of leaky females, (E) male flower biomass and (F) plant aboveground biomass. The prediction intervals evaluate whether the replicate study estimates differ from the original estimates more than expected only due to sampling error. Ctr: control treatment, Sel: selection treatment, G1: generation 1, G3: generation 3.

Despite the methodological differences between both studies, the mean effects of treatment and generation on the traits under selection (female leakiness and male flower production) in the replicate did not differ substantially from measurements made in the original study more than expected due to sampling error (Figure 2A-C). We predicted sex allocation metrics would be sensitive to our methodological differences, which underestimate female sex allocation by around 30% (Villamil et al., 2021) and male sex allocation by around 10%, and expected them to fall outside the prediction intervals – given that the variation was no longer due only to sampling error (Spence and Stanley, 2016; Schauer and Hedges, 2021). However, the mean effects of generation and selection on female leakiness and male flower production actually fell within the prediction intervals of the original study (i.e., were successfully replicated).

**Figure 2.**
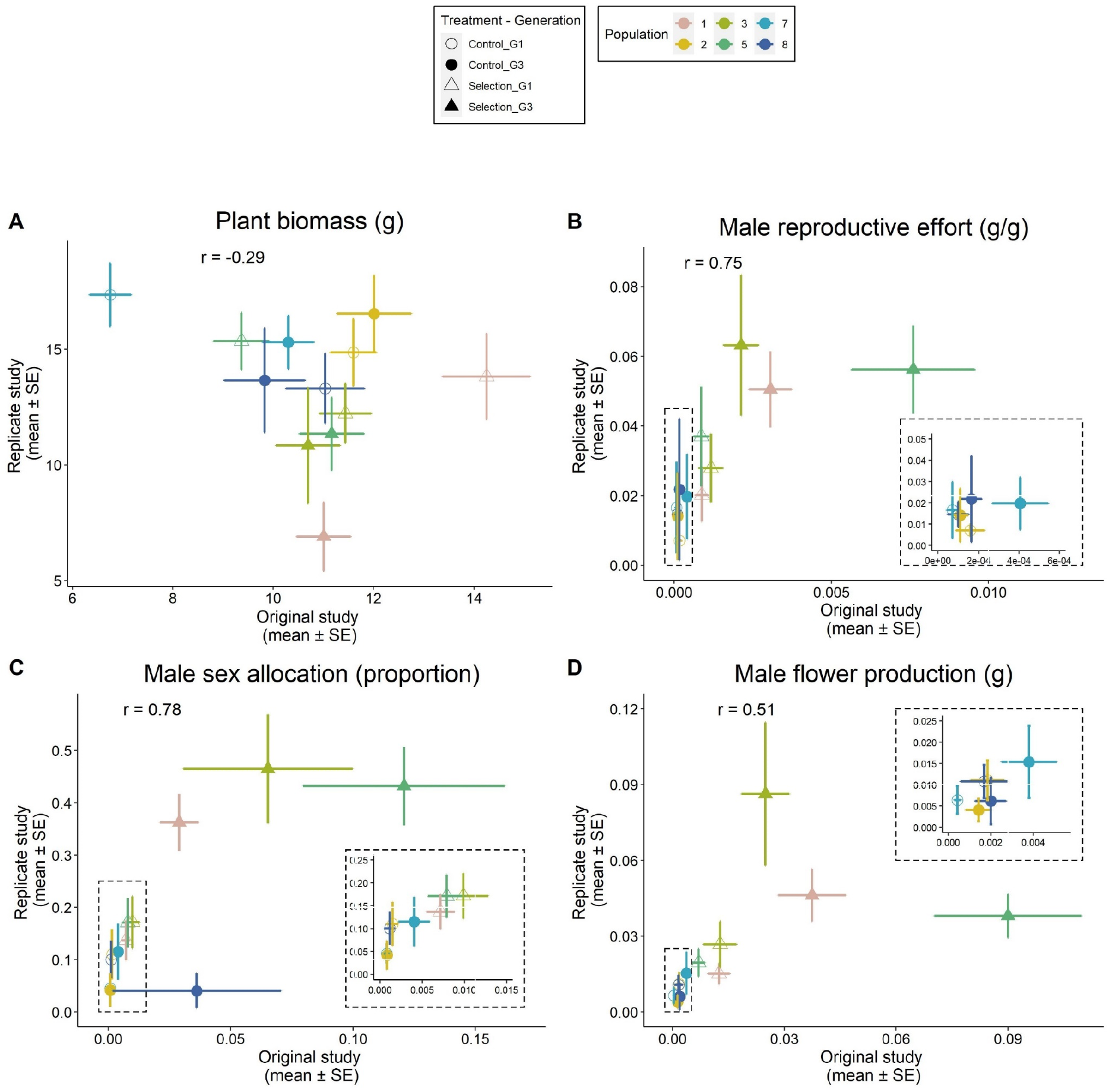
Between and within population variation correlating estimates from the original study (2016) *versus* the replicate study (2020) for all individual populations from the two generations that were replicated (G1, G3). (A) aboveground plant biomass, (B) male reproductive effort defined as the grams of male flowers produced per gram of plant biomass, (C) male sex allocation defined as the proportion of male flowers per gram of reproductive biomass, and (D) net male flower production. The dashed squares contain a zoomed in section of the plot.

In line with our initial prediction, the mean values of female leakiness and male flower production per treatment group differed from the original more than expected due to sampling error. Only four of the 12 replicates, mean values fell within the prediction intervals (Figure 2D-F). Females in the replicate study were on average 30% less leaky compared with the counterparts from the original study. Nevertheless, selected females were consistently leakier than controls, and G3 females were leakier than G1 females (Figure 2D). These difference may be due to a reduced sample size in the replicate, or may reflect the expected underestimation of male flowers derived from the subsampling procedure. Regarding total male flower production, we observed the opposite trend, where females in the replicate produced on average 70% more male flowers than those in the original study (Figure 2E).

Such an increase is likely due to the greater age of plants in the replicate study, because older plants are larger, and plant size influences resource availability and reproductive investment in *M. annua* (Tonnabel et al., 2019) and many other plant species (Charnov and Bull, 1977; Freeman et al., 1981; Charnov, 1982; de Jong and Klinkhamer, 1989; Klinkhamer et al., 1997). Taken together, these two results suggest that our subsampling procedure may be missing a third of the leaky females with very low male flower production – possibly those that only produced male flowers in basal branches that were excluded by our subsampling procedure. Overall, the successful replication of the effect size magnitudes and directions of treatment and generation on female leaky sex expression, despite methodological variation in sampling and culturing, indicates that the evolution of enhanced leakiness in females of *M. annua* in response to mate limitation is a strongly generalizable (sensu Fraser et al., 2020) biological pattern, robust to methodological and environmental changes.

### Repeatability of patterns of among-population variation

We observed considerable among-population variation in the measured traits, as also reported by Cossard et al. (2021a). In both studies, while population identity explained a negligible amount of variance (< 0.001%) in traits under selection (leakiness and male flower production) (Table 1), it explained substantial variation in plant biomass (range: 0.58-17.05 %). The fact that this result was not entirely consistent between studies is likely due to the environmental and culturing differences between the original and replicate studies (Table 1). When plotting and estimating the correlation between population mean values for both common gardens, we observed that, for all sex-allocation traits measured, females clustered according to their treatment and generation group, indicating that the overall patterns of among-population variation were fairly consistent in both studies. Females in the treatment populations from generation G3 had the highest male sex allocation values for all three traits and clustered at the top right corner, whereas those from generation G1 clustered together closer to the control groups, which aggregate at the bottom left corner (Figure 2B-D). In contrast, plant biomass showed no clustering by treatment groups, as expected.

Among-population differences in the treatment group may be the result of pre-existing variation in male flower production, e.g., due to difference in the presence of specific alleles, or in their frequency, among replicate populations at the time the experiment was first established. The fact that population 5 showed the greatest male flower production in both common garden studies in G3 would be consistent with this sort of genetic effect. In more general terms, the difference among replicates recorded by Cossard et al. (2021a) were similar and largely reproduced in our common garden, with a significant correlation between them. Apart from pointing to likely genetic differences among populations within treatment groups, this result confirms that male sex allocation and male reproductive effort are relatively robust to the sort of environmental differences experienced by the plants between experiments, likely because they are relative ratios that account for the substantial plant size differences observed between studies: male sex allocation showed the strongest correspondence between our two common gardens, followed closely by male reproductive effort. These results contrast with results for male flower biomass, which showed a ∼25% weaker correlation (Figure 2B-D). Again, among-population variation in plant above-ground biomass estimates was not correlated between both studies (Figure 1A).

## Conclusions

Our study confirmed the strong treatment effects reported by Cossard et al.(2021b), but its more interesting result is the confirmation, through successful replication, of much of the among-population variation reported by Cossard et al. (2021b). The two common gardens were conducted in different years and under different weather conditions. While traits were measured for relatively young plants by Cossard et al. (2021b), here we measured much older, almost senescent individuals. The similarities in the relative value of traits for specific populations between the two common gardens are thus noteworthy and suggest that at least a substantial element of the among-population variation reported by Cossard et al. (2021b) can be attributed to genetic differences among them. These differences may be the result of sampling differences at the beginning of the selection experiment, but they may also be the result of unmeasured differences in the growth conditions experienced among populations over the course of their evolution. Detailed analysis of genomic variation among the replicate populations will help further to establish the nature of their divergence.

## ACKNOWLEDGEMENTS

We thank Ehouarn Le Faou for comments on the manuscript, Aline Revel for assistance in growing plants, a team of student assistants for help with phenotyping plants, and the Swiss National Science Foundation (grant 310030_185196 to JRP) and the University of Lausanne for funding.

## Authors contributions

JRP and NV conceived the idea, NV and XL collected the data, NV analysed the data, NV and JRP wrote the paper.

## Data availability statement

The data and code used to generate the results will be uploaded to the XXXX repository available at [link].

## SUPPLEMENTARY MATERIALS

### Study system

*Mercurialis annua* (Euphorbiaceae) is an annual wind-pollinated herb distributed throughout central and western Europe and around the Mediterranean Basin (Tutin et al., 1968). The species has long been used as a model system to investigate the evolution of dioecy and sex expression (Yampolsky, 1919; Yampolsky, 1930; Pannell, 1997; Obbard et al., 2006; Pannell et al., 2008) due to its great diversity and plasticity in sex expression and sexual systems (Yampolsky, 1930; Pannell, 1997; Cossard and Pannell, 2019; Cossard et al., 2021a). *M. annua* has chromosomal sex determination (XX♀; XY♂) (Russell and Pannell, 2015; Veltsos et al., 2018; Veltsos et al., 2019), mediated by endogenous hormonal signalling (Durand and Durand, 1991).

### Plant culturing

Plants were sowed in trays with sterilised soil (163 Ricoter) and germinated under greenhouse conditions at the University of Lausanne, Switzerland, with controlled temperature (25^°^C) and humidity (50%). In the original study, plants were sexed and potted after 6 weeks, whilst in the replicate study plants stayed in the germination trays for 10-12 weeks due to restricted access to University facilities during the 2020 COVID-lockdown.

In both studies, adult plants were grown in individual pots (Teku series TO 14D) with soil (140 Ricoter soil) and slow-release fertiliser (Hauer-Tardit 3M: 500 g per 100 L of soil). Plants were kept within a polytunnel with automatic watering under semi-controlled temperature conditions, regulated by the automatic opening of the polytunnel ventilation windows when temperature reached 14°C, and closing them at temperatures below 14^°^C. In both experiments, plants were sown in spring and grown over the summer months.

Plants in the replicate study suffered damage from thrips and viral infections and experienced a brief period of drought due to a failure in the automatic watering system over a weekend. Plants were allowed to recover fully before being measured, but we lost some individuals due to disease and drought. In the original study, plants were harvested 14 weeks after germination, whilst plants in the replicate study were harvested 20 weeks after germination - again as a consequence of access restrictions during the 2020 COVID-lockdown.

### Sampling methods

Plant sampling consisted of cutting all above-ground plant material, recording total height and phenotyping the plant. Phenotyping consisted of carefully examining the plant, counting the number of fruits present, and harvesting all male flowers using tweezers. Male flowers were stored in paper envelopes, dried and then weighed. After phenotyping, plants were placed in individually labelled plastic bread bags, dried and then weighed. To estimate seed production, seeds were isolated from the dried plant material, stored in paper envelopes and weighed. All materials were dried in an oven at 50^°^C for at least 14 days and weighed using a digital scale.

In the original study, the whole plant was phenotyped. In the replicate study we subsampled the plant by cutting the top 30 cm of the plant and phenotyped only this segment. Both plant segments were individually labelled, dried and weighed. In the replicate, seeds were only isolated from the apical dried plant segment (subsample).

